# Natural variation in soybean perception of bacterial quorum-sensing signals shapes primed defense

**DOI:** 10.64898/2026.08.14.744918

**Authors:** Shimaa Adss, Ahmed Elhady, Ludger Beerhues, Volker Hahn, Holger Heuer

## Abstract

Plants perceive N-acyl homoserine lactones (AHLs), the quorum-sensing signals of rhizobacteria, and can translate them into a primed state of enhanced inducible immunity. Whether this perception-to-defense translation varies heritably among genotypes is largely unknown. Using soybean (Glycine max) responses to N-3-oxo-tetradecanoyl-L-homoserine lactone (oxo-C14-HSL), with root invasion by the nematode Pratylenchus penetrans as a quantitative challenge revealing the primed state, we tested whether AHL-triggered priming is genotype-dependent. Priming with the oxo-C14-HSL-producing rhizobacterium Ensifer meliloti ExpR+ potentiated defense in the responsive cultivar Primus, reducing nematode invasion (P < 0.001), but not in the weakly responsive Sigalia (P = 0.082). The purified molecule reproduced this contrast, localizing the difference to signal transduction within the plant rather than bacterial colonization. Correspondingly, the phytoalexin glyceollin accumulated in a priming- and challenge-dependent manner in Primus but not Sigalia, the formal signature of priming. Across a Sigalia × Primus recombinant inbred line population and an independent diversity panel, responsiveness was heritable, continuously distributed, and transgressive. Two genome-wide association studies and a cross-population meta-analysis converged on a well-calibrated null, indicating a polygenic or strongly environment-dependent architecture rather than a common-variant, large-effect-size model. Translating a bacterial signal into primed defense is thus a quantitatively varying plant trait.

**Highlight:** Soybean genotypes differ heritably in their ability to convert a bacterial quorum-sensing signal into primed phytoalexin defense, revealing this translation step as a quantitative, polygenic trait.

## 1. Introduction

A central feature of plant immunity is its inducibility: rather than maintaining costly defenses constitutively, plants sense cues from their biotic environment and adjust the amplitude and speed of their defense responses accordingly. Among the most consequential of these cues are signals exchanged with the root-associated microbiota. Many Gram-negative rhizobacteria communicate through N-acyl homoserine lactones (AHLs), quorum-sensing molecules that coordinate population-level behavior (Fuqua et al., 2001), and plants have evolved the capacity to perceive these bacterial signals and mount adaptive responses (Ortíz-Castro et al., 2008; Hartmann et al., 2014). How reliably an individual genotype converts such a perceived signal into effective, inducible defense, and whether that capacity varies heritably within a crop species, is the question we address here.

Defense priming provides a powerful framework for this question. A priming stimulus does not itself activate Defenses; instead, it sensitizes the plant so that, upon subsequent challenge, Defenses are deployed faster and more strongly, at little fitness cost in the absence of attack (van Hulten et al., 2006; Conrath et al., 2015; Martínez-Medina et al., 2016; Mauch-Mani et al., 2017). Beneficial soil microorganisms establish primed states, such as induced systemic resistance (Wees et al., 2008; Pieterse et al., 2014), and long-chain AHLs, notably N-3-oxo-tetradecanoyl-L-homoserine lactone (oxo-C14-HSL), are potent priming cues (Schikora et al., 2011; Schenk et al., 2014). Mechanistically, AHL-priming in Arabidopsis and barley operates through augmented and prolonged activation of the mitogen-activated protein kinases MPK3 and MPK6, potentiated defense-gene expression, and reinforcement of the salicylic acid and oxylipin pathways, culminating in cell-wall fortification (Schikora et al., 2011; Schenk et al., 2014; Schenk and Schikora, 2014). Because priming is defined by a potentiated response to challenge, it requires a quantitative Defense output and a defined challenge to be measured. In soybean (Glycine max (L.) Merr.), earlier work by our group established such a system: priming cv. Primus with oxo-C14-HSL or the oxo-C14-HSL-producing rhizobacterium Ensifer meliloti ExpR+ potentiated accumulation of the phytoalexin glyceollin and reduced root invasion by the migratory nematode Pratylenchus penetrans (Adss et al., 2021). Root-lesion nematodes, which feed destructively within the root cortex and cause widely underestimated yield losses in soybean and other crops as chemical controls are withdrawn (Castillo and Vovlas, 2007; Jones and Fosu-Nyarko, 2014; Elhady et al., 2018; Mokrini et al., 2019; Oka, 2020), thus provide a stringent, quantitative challenge with which to read out the primed state.

Crucially, the capacity to respond to AHL signals is not universal. In barley, genetic differences among accessions govern responsiveness to oxo-C14-HSL (Shrestha et al., 2019), and AHL-priming efficiency varies among cereal genotypes (Hernández-Reyes et al., 2014; Shrestha et al., 2022). Plants also differ in their responses to distinct AHL molecules, indicating specificity in perception and downstream signaling (Ortíz-Castro et al., 2008; Schenk and Schikora, 2014). If such genotype dependence operates in soybean, the ability to translate a perceived AHL signal into inducible Defense would itself be a heritable, quantitatively varying trait, amenable to genetic dissection. This possibility has previously not been tested in soybean and is the central question of this study.

Glyceollins are prenylated pterocarpan phytoalexins produced through the legume-specific isoflavonoid branch of the phenylpropanoid pathway and are central to soybean Defense against diverse pathogens (Lygin et al., 2013; Jahan and Kovinich, 2019; Jahan et al., 2020). Their biosynthesis is inducible and tightly regulated, requiring multiple enzymatic steps from daidzein, and is controlled by a network of transcriptional activators and repressors (Graham et al., 2007; Akashi et al., 2009; Lin et al., 2024; Xie et al., 2025). This regulatory architecture provides plausible molecular substrates on which genotype-dependent differences in priming responsiveness could act.

We tested the hypothesis that the capacity of soybean to translate the bacterial signal oxo-C14-HSL into primed Defense is genotype-dependent, and that variation in inducible glyceollin accumulation, rather than in signal perception per se, underlies contrasting responsiveness. We further sought to define the genetic architecture of this trait. To this end, we (i) quantified the primed Defense response, as inducible glyceollin accumulation and, as a stringent functional read-out, suppression of nematode invasion, in the contrasting cultivars Primus and Sigalia following exposure to live E. meliloti ExpR+ and to purified oxo-C14-HSL; (ii) phenotyped priming-induced glyceollin accumulation across a Sigalia × Primus recombinant inbred line (RIL) population and an independent 203-accession diversity panel; and (iii) dissected the genetic basis through three complementary, power-aware analyses: a RIL genome-wide association study (GWAS), an independent diversity-panel GWAS built on public SoySNP50K genotypes, and a cross-population meta-analysis. Together, this design treats the primed phytoalexin response as a quantitative trait, rigorously characterizes its genetic architecture, and tests the robustness of the conclusions across populations and marker platforms.

## 2. Materials and Methods

### 2.1 Plant material and soybean populations

Two soybean cultivars with contrasting priming behavior served as references throughout: cv. Primus (highly responsive) and cv. Sigalia (weakly responsive) (Adss et al., 2021). For genetic analysis, RIL population derived from the cross Sigalia × Primus (F7) was used together with the parental lines and the reference cultivar Protina. An independent panel of 203 genetically diverse soybean accessions, comprising USDA Plant Introduction (PI) accessions and registered European and Asian cultivars, was assembled to assess the breadth of priming responsiveness across germplasm (Hahn and Würschum, 2014). Seed material and accompanying passport data for the RIL population, its parents, and the diversity panel were provided by Prof. Volker Hahn, State Plant Breeding Institute, University of Hohenheim, Germany. The RIL population carries the breeding designation SJE09-002-01. Accession identities and pedigrees for the diversity panel were curated from the project genotype registry. The complete list of accessions, including their USDA Plant Introduction numbers and public-genotype availability, is provided in Supplementary Table S2.

### 2.2 Bacterial strains and priming treatments

The oxo-C14-HSL-producing strain Ensifer meliloti ExpR+ and the isogenic AHL-degrading strain E. meliloti attM (expressing the AttM lactonase) served as priming-competent and priming-deficient controls, respectively, as described previously (Hernández-Reyes et al., 2014; Adss et al., 2021). Bacteria were grown to the required density and applied as a root drench, and the purified quorum-sensing molecule oxo-C14-HSL (Sigma-Aldrich) was applied following the protocol of Shrestha et al. (2022) to distinguish signal perception from other bacterial traits. Control plants received the corresponding bacterium-free or solvent-only treatment (Hernández-Reyes et al., 2014; Adss et al., 2021; Shrestha et al., 2022).

### 2.3 Nematode culture and inoculation bioassays

Pratylenchus penetrans was cultured, and mixed-stage nematodes were extracted for inoculation following the procedures of Adss et al. (2021). Seedlings of Primus and Sigalia were primed with E. meliloti ExpR+, E. meliloti attM, or purified oxo-C14-HSL, or received the control, and were subsequently challenge-inoculated with P. penetrans. Invaded nematodes were stained and counted at 10 days post-infection, and density was expressed as nematodes per plant (Adss et al., 2021). Because counts were right-skewed, values were log10-transformed prior to analysis, with log values computed directly from raw counts.

### 2.4 Glyceollin quantification by cotyledon assay

Priming-induced glyceollin accumulation was quantified following the phytoalexin analysis procedure of Adss et al. (2021). Detached cotyledons from primed (oxo-C14-HSL; P) or non-primed (C) seedlings were treated and, for the mechanistic experiment, additionally challenged with *P. penetrans* extract, so that every cotyledon received nematode extract and the priming contrast (P minus C) reflected the true priming effect rather than a difference in elicitation. Glyceollin was extracted and quantified spectrophotometrically; absorbance was normalized to cotyledon fresh weight and expressed as optical density per gram (OD g⁻¹). For each genotype, priming responsiveness was quantified as the priming effect (P minus C) and, secondarily, as the fold induction (P/C).

### 2.5 Phenotyping of the RIL population and the diversity panel

The RIL population and the 203-accession diversity panel were each phenotyped for priming-induced glyceollin accumulation using the cotyledon assay. The RIL population was evaluated in two independent experiments, and the diversity panel in three, each with multiple replicate cotyledons per genotype and treatment. Treatment means were computed per genotype across replicates and experiments, and the priming effect (P minus C) and fold induction were derived. For the diversity panel, absorbance was recomputed from raw optical density and cotyledon weight measurements to ensure that the derived values reflected the corrected primary data. Cross-experiment repeatability of the priming effect was estimated as the Pearson correlation of per-genotype priming values between independent experiments.

### 2.6 Genotyping and marker quality control

The Sigalia × Primus RIL population was genotyped by genotyping-by-sequencing (GBS) following the ApeKI-based protocol of Elshire et al. (2011), and SNPs were called and processed into a HapMap genotype matrix with the TASSEL-GBS pipeline (Glaubitz et al., 2014), anchored to the Glycine max Wm82.a2 reference assembly (Schmutz et al., 2010). Phenotype and marker records were reconciled through the GBS key file by pedigree, matching each phenotyped line to its genotype. From 98,309 raw SNPs, markers were filtered to remove monomorphic sites, sites with more than 20% missing calls, and sites with minor allele frequency (MAF) below 0.05; one line with more than 99% missing data was excluded, yielding 9,573 SNPs across 80 RILs. For the diversity panel, genome-wide genotypes for the USDA PI accessions were obtained from the public SoySNP50K resource (Illumina Infinium BeadChip; 42,291 SNPs) in the Wm82.a2 coordinate system (Schmutz et al., 2010; Song et al., 2013). Allele calls were converted to numeric dosage; after removal of monomorphic markers and filtering for missingness at most 20% and MAF at least 0.05, 33,300 SNPs remained across 135 accessions that had both genotype and phenotype data. Missing genotypes were imputed to the marker mean for downstream analyses.

### 2.7 Population structure, kinship, and genome-wide association analysis

For each population, population structure was examined by principal component analysis (PCA) of the marker matrix (Price et al., 2006), and genome-wide relatedness was estimated as a realized additive (VanRaden) kinship matrix (Yu et al., 2006). Association between the priming effect (P minus C; primary trait) and each SNP was tested with a mixed linear model that accounted for kinship and leading principal components (Q + K) (Price et al., 2006; Kang et al., 2010), implemented through rrBLUP (Kang et al., 2010) and an efficient variance-component (EMMAX-type) procedure for the larger diversity-panel marker set (Price et al., 2006). Fold induction (log2 P/C) was analyzed as a secondary trait, and a general linear model with principal-component covariates was fitted in parallel as a robustness check for the RIL population. Because standard corrections are conservative under strong linkage disequilibrium and limited sample size (Churchill and Doerge, 1994; Huang et al., 2019), an empirical genome-wide significance threshold was derived by permutation (1,000 shuffles of the phenotype residuals; 95th and 90th percentiles of the maximum -log10 P) (Korte and Farlow, 2013). Genomic inflation was summarized by the genomic control factor (lambda_GC) (Benjamini and Hochberg, 1995) and quantile-quantile plots. Candidate genes within the linkage-disequilibrium-defined support intervals of leading associations were retrieved from the Wm82.a2 annotation (Cao et al., 1997; Subramanian et al., 2005) and prioritized by documented roles in isoflavonoid and glyceollin biosynthesis, phytoalexin regulation, and systemic-resistance signaling (Akashi et al., 2009; Withers and Dong, 2016); allelic effects at leading SNPs were examined by grouping RILs by parental (Primus-type versus Sigalia-type) allele.

### 2.8 Cross-population meta-analysis

Because the RIL (GBS) and diversity-panel (SoySNP50K) datasets were genotyped on different platforms with few shared SNPs, the two GWAS were combined at the level of genomic regions rather than by pooling individuals (Kotz, 1992). The genome was partitioned into 500-kb windows; within each window, the strongest signal (minimum p) was retained per population. For windows genotyped in both populations, per-window p-values were combined by Fisher’s method (Chang et al., 2016), and false-discovery-rate correction was applied across shared windows. Genome-wide concordance of window-level signals between populations was assessed by Pearson correlation, and the RIL candidate regions were examined directly for support in the diversity panel.

### 2.9 Statistical analysis and data visualization

For two-treatment bioassays, priming effects were tested within each genotype by two-sample t-tests. For the three-treatment (attM/ExpR+/control) and four-treatment glyceollin experiments, groups were compared within genotype by one-way analysis of variance followed by Tukey’s honestly significant difference test, with significance shown as compact-letter displays; differences were considered significant at P < 0.05. All analyses and figures were produced in R version 4.3.3 (Hahn and Würschum, 2014) using ggplot2 (v3.4.4), with genome-wide association analyses implemented in rrBLUP (v4.6.3) (Kang et al., 2010) and the cross-population meta-analysis performed in Python 3.12 using SciPy and statsmodels. Boxplots show the median, interquartile range, and 1.5 times the interquartile range whiskers, with significance letters placed above the upper whisker. Figures were exported at 600 dpi. Diversity-panel genotypes are publicly available from SoyBase (SoySNP50K) (Pandey and Somssich, 2009).

## 3. Results

### 3.1 The primed Defense response to oxo-C14-HSL is genotype-dependent

Priming with the oxo-C14-HSL-producing rhizobacterium E. meliloti ExpR+ reduced P. penetrans invasion in a genotype-dependent manner (Figure 1A). In the responsive cultivar Primus, ExpR+ priming markedly reduced root nematode density relative to the control (P < 0.001), whereas in the weakly responsive cultivar Sigalia the reduction was not significant (P = 0.082). The purified oxo-C14-HSL molecule reproduced this contrast (Figure 1B): Primus again showed a significant reduction (P = 0.013) while Sigalia did not (P = 0.84). Because the pure signal recapitulated the response elicited by the live bacterium, the genotype-dependent difference resides in the plant response to the AHL signal rather than in the bacterial colonization, consistent with the specificity of AHL perception reported in other species (Ortíz-Castro et al., 2008; Schenk and Schikora, 2014; Shrestha et al., 2019). Restricting the comparison in Sigalia to the AHL-producing (ExpR+) versus AHL-degrading (attM) strains produced no significant differences (Supplementary Fig. S1), consistent with the limited capacity of Sigalia to translate the signal into effective Defense. Soybean cultivars, including Primus, differ in their basal susceptibility to Pratylenchus under temperate conditions (Elhady et al., 2019).

**Figure 1.**
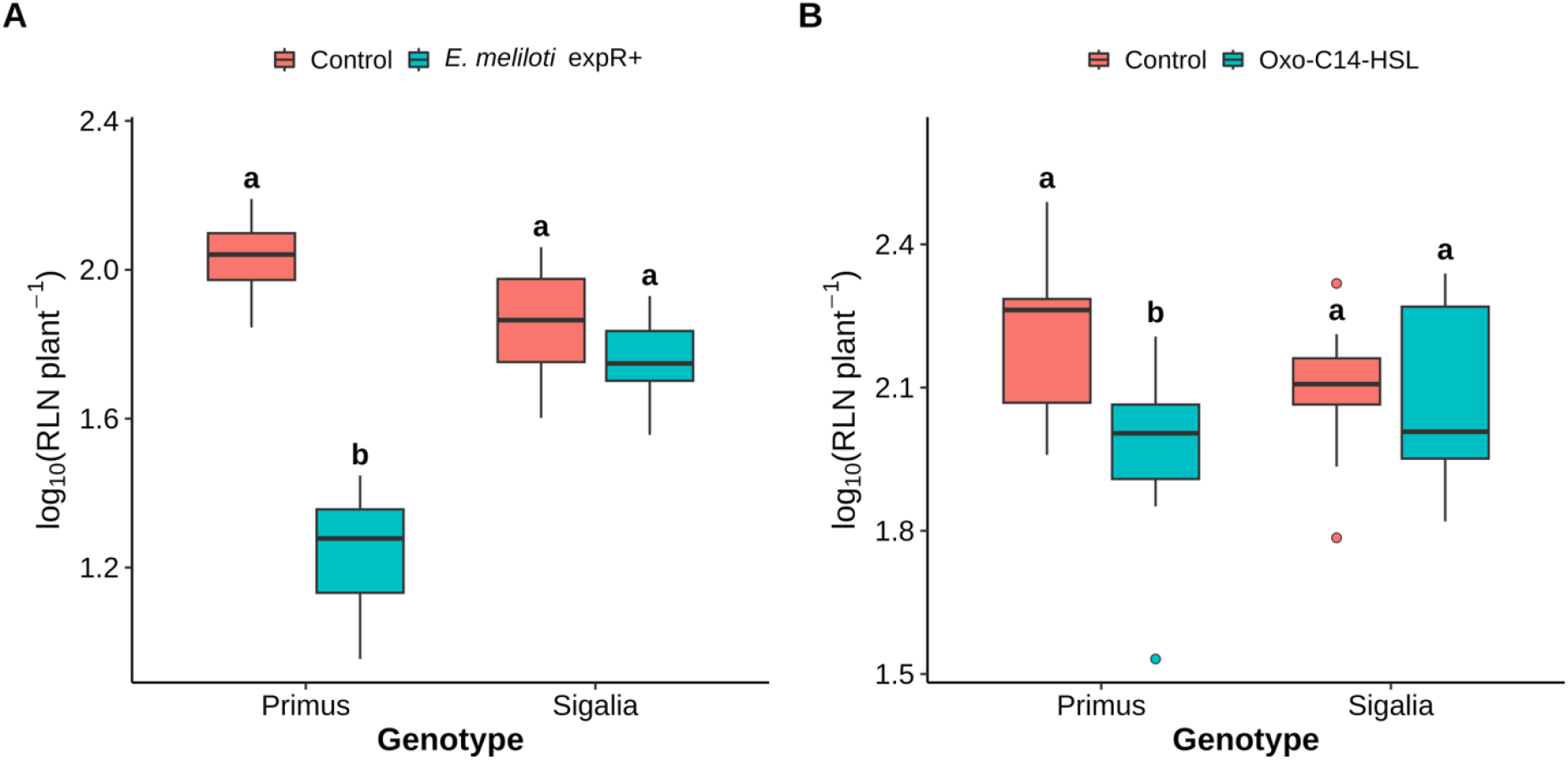
Genotype-dependent suppression of *Pratylenchus penetrans* by AHL priming. Root nematode density (log10 nematodes per plant) in Primus and Sigalia after priming with (A) *Ensifer meliloti* ExpR+ or (B) purified oxo-C14-HSL, relative to controls. Boxes show the median and interquartile range; whiskers extend to 1.5 times the interquartile range. Within each genotype, different letters denote a significant difference between treatments (t-test, P < 0.05). Fig. 1 alt text. Grouped bar chart comparing root nematode density on a logarithmic scale in soybean cultivars Primus and Sigalia. Panel A shows priming with Ensifer meliloti ExpR+ and panel B priming with purified oxo-C14-HSL, each against its control. Nematode density falls markedly in Primus but changes little in Sigalia.

### 3.2 Priming induces glyceollin accumulation only in the responsive genotype upon challenge

To test whether glyceollin underlies the genotype-dependent protection, its accumulation was quantified across four treatments (control, P. penetrans challenge, oxo-C14-HSL priming, priming plus challenge; Figure 2). In Primus, glyceollin accumulated strongly and significantly only when priming was combined with nematode challenge, not in response to priming or challenge alone (ANOVA P < 0.001). In Sigalia, glyceollin remained low across all treatments and showed no coordinated priming-plus-challenge induction. This is the defining signature of priming: the responsive genotype is sensitized to produce the phytoalexin rapidly upon attack rather than accumulating it constitutively, minimizing metabolic cost (van Hulten et al., 2006; Conrath et al., 2015; Mauch-Mani et al., 2017).

**Figure 2.**
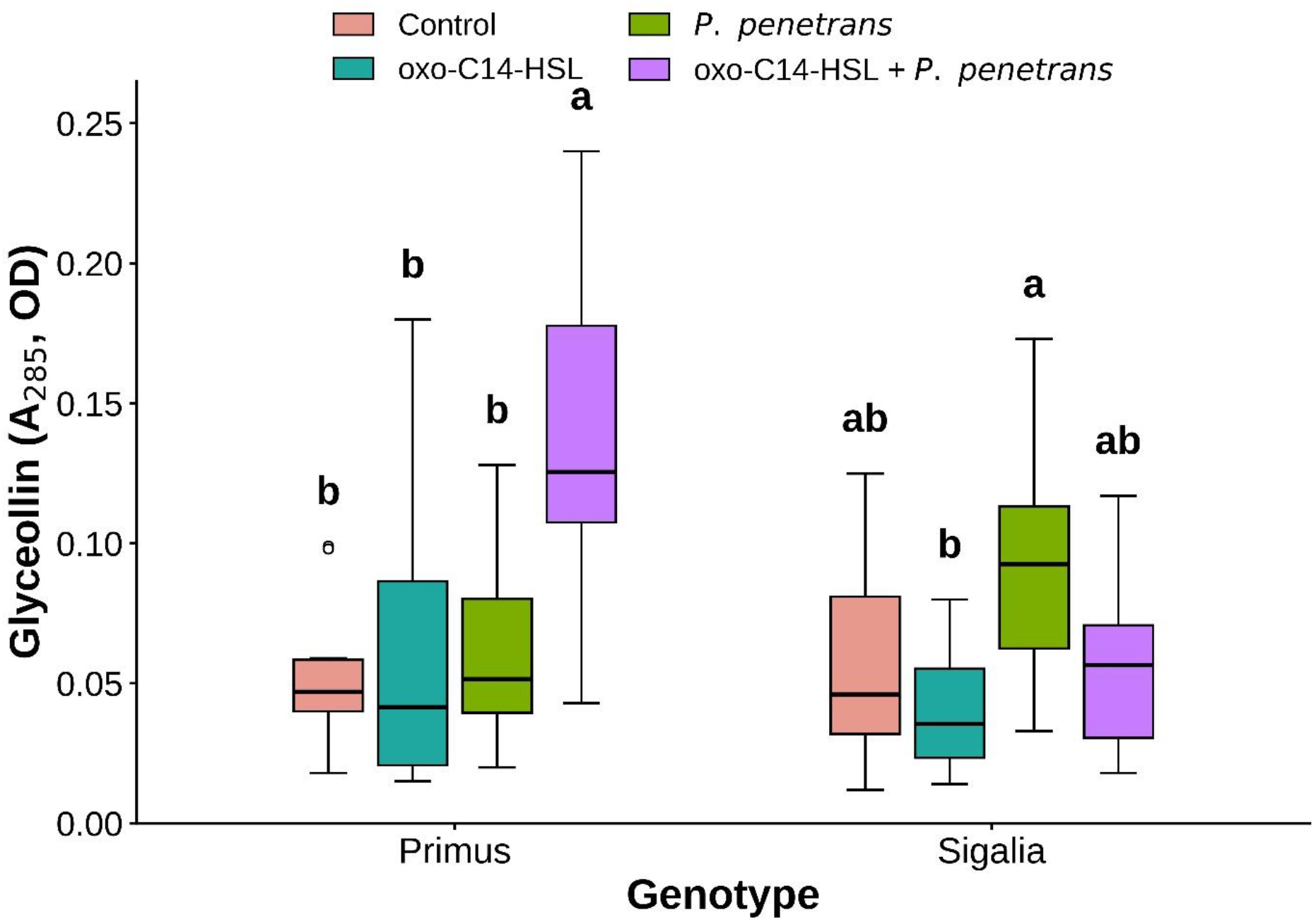
Priming-induced glyceollin accumulation in Primus and Sigalia. Glyceollin content (absorbance, OD) in cotyledons under control, *P. penetrans* challenge, oxo-C14-HSL priming, and priming plus challenge. Within each genotype, different letters denote significant differences among treatments (Tukey HSD, P < 0.05). Fig. 2 alt text. Box plots of glyceollin absorbance in cotyledons of Primus and Sigalia across four treatments: control, nematode challenge alone, oxo-C14-HSL priming alone, and priming followed by challenge. Glyceollin rises sharply only in primed and challenged Primus; Sigalia shows no comparable rise.

### 3.3 Priming responsiveness is heritable and continuously distributed across two independent populations

To determine whether the Primus and Sigalia contrast reflects broader genetic variation, priming-induced glyceollin accumulation was quantified across the Sigalia × Primus RIL population. The priming effect (P minus C) varied continuously from strongly positive to negative (Figure 3). The responsive parent Primus ranked among the most responsive lines (priming effect +0.285 OD g⁻¹; rank 3 of 85), the weakly responsive parent Sigalia fell near the middle with an essentially neutral response (- 0.018 OD g⁻¹), and the reference cultivar Protina was mildly negative (-0.072 OD g⁻¹). Transgressive segregation beyond both parents indicated that the trait was quantitative and influenced by multiple loci; the same pattern was observed for fold induction (Supplementary Figures S2 and S3).

**Figure 3.**
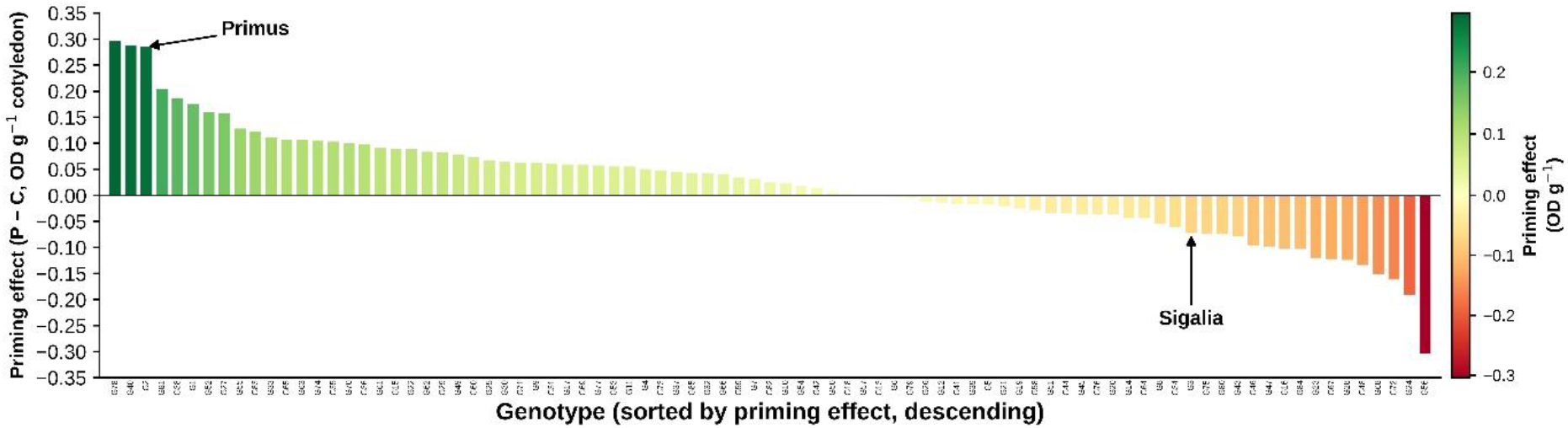
Continuous variation in priming responsiveness across the Sigalia × Primus RIL population. Bars show the priming effect (P minus C, OD g⁻¹ cotyledon) per line, ordered from most to least responsive and colored on a red-to-green scale. Parents Primus and Sigalia and the reference cultivar Protina are labeled. Fig. 3 alt text. Waterfall bar chart of the priming effect for each recombinant inbred line of the Sigalia by Primus population, ordered from the most positive to the most negative response. Bars are coloured on a continuous scale and span positive and negative values, with the parental cultivars indicated.

An independent panel of 203 diverse accessions confirmed that priming responsiveness is widespread and continuously distributed (Figure 4). The priming effect ranged from strongly negative to strongly positive, with approximately half of the accessions showing positive priming. The responsive cultivar Primus (accession 3327) ranked near the top of the distribution (rank 5 of 203), consistent with its reference status, whereas Sigalia (accession 3363) occupied an intermediate rank; numerous accessions exceeded Primus, revealing favorable alleles dispersed across germplasm (Hahn and Würschum, 2014). Because the two populations differ in genetic background, marker platform, and experimental context, their concordant demonstration of heritable, continuous variation provides robust evidence that AHL-priming responsiveness is a genuine quantitative trait.

**Figure 4.**
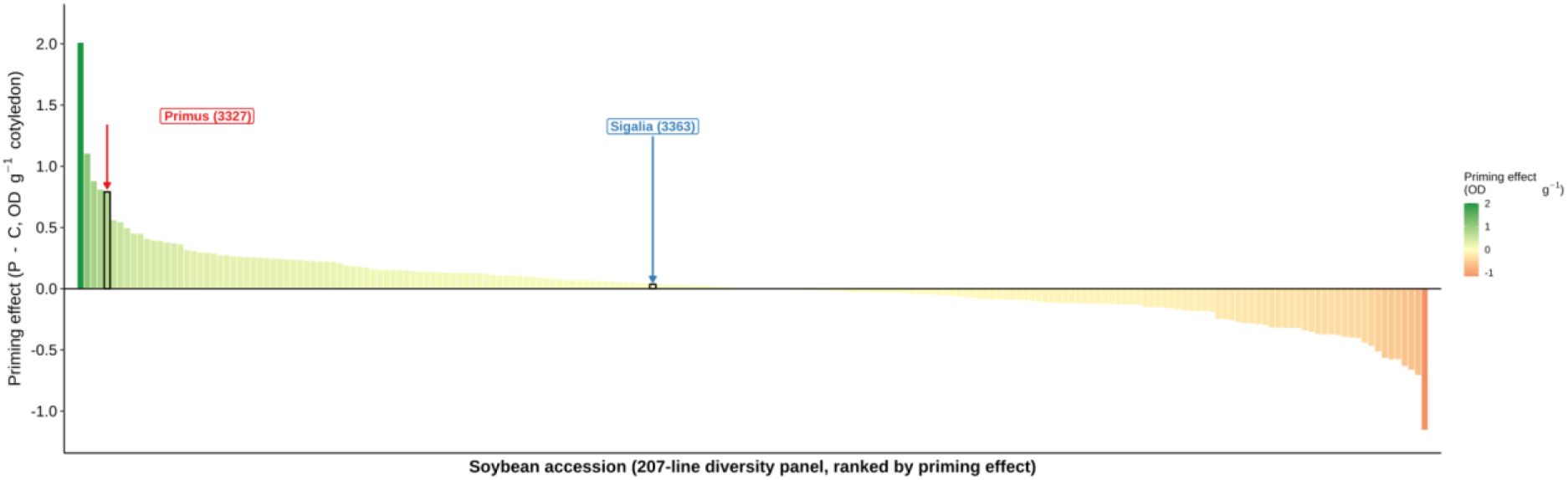
Priming responsiveness across the independent diversity panel of 203 soybean accessions. Bars show the priming effect (P minus C, OD g⁻¹ cotyledon) per accession, ranked and colored on a red-to-green scale. Primus (3327) and Sigalia (3363) are marked with arrows and their registry codes; many accessions exceed the responsiveness of the reference cultivar, Primus. Fig. 4 alt text. Waterfall bar chart of the priming effect for each accession of the independent diversity panel, ranked from most to least responsive. The distribution is continuous, with many accessions exceeding the responsive parent.

### 3.4 Priming responsiveness shows no common variant of large effect across three complementary analyses

RIL population. The priming effect was analyzed by GWAS in the RIL population (80 lines, 9,573 SNPs). Principal component analysis showed no strong stratification, and linkage disequilibrium decayed to half-maximum over approximately 1.8 Mb, reflecting the biparental origin of the population (Supplementary Figures S4, S5). The mixed linear model was well-behaved (lambda_GC approximately 0.92; Figure 5). No SNP surpassed the permutation-derived genome-wide significance threshold (95th-percentile -log10 P approximately 4.65; observed maximum approximately 3.40), a power-aware null consistent with the modest population size and with the low cross-experiment repeatability of the phenotype (Pearson r approximately 0.17). The general linear model and the fold-induction analysis yielded concordant results (Supplementary Figures S7 and S9).

**Figure 5.**
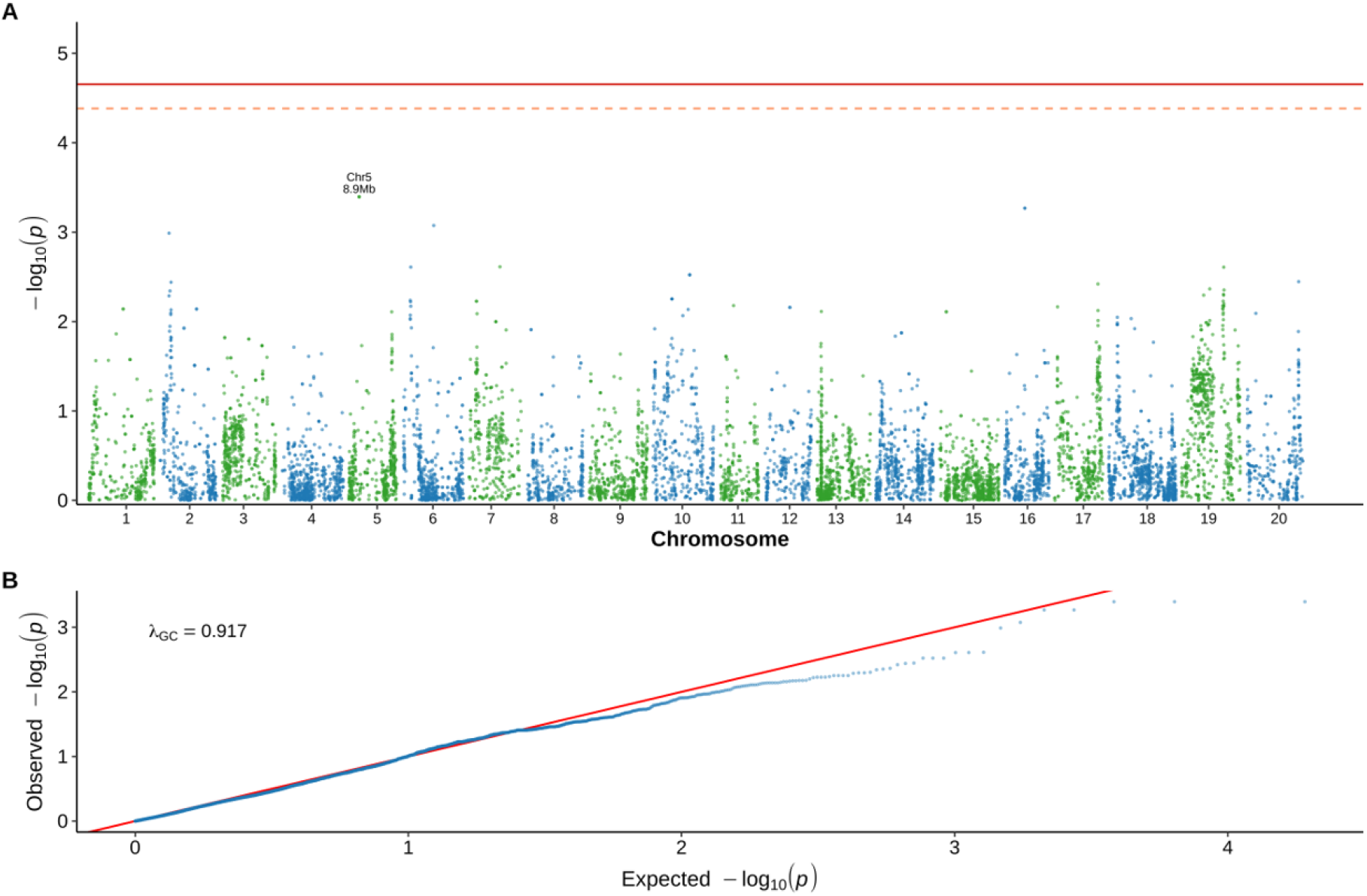
Genome-wide association analysis of the priming effect in the Sigalia × Primus RIL population. (A) Manhattan plot of -log10 P from the mixed linear model across the 20 chromosomes; solid and dashed lines denote the permutation-derived genome-wide significant (95th percentile) and suggestive (90th percentile) thresholds. (B) Quantile-quantile plot with the genomic control factor (lambda_GC). Fig. 5 alt text. Genome-wide association results for the recombinant inbred line population. Panel A is a Manhattan plot of negative log-transformed P values across the twenty chromosomes with the permutation threshold marked; no marker exceeds it. Panel B is a quantile-quantile plot lying close to the expected diagonal.

Diversity panel. An independent GWAS in the diversity panel (135 accessions, 33,300 SoySNP50K markers) provided a stronger, orthogonal test with denser markers on an unrelated set of accessions (Song et al., 2013). The model was exceptionally well-calibrated (lambda_GC = 0.99; Figure 6), yet again no marker approached the permutation threshold (95th percentile approximately 5.10; observed maximum approximately 3.79; minimum false-discovery rate approximately 0.99). The strongest, though non-significant, signals formed a small cluster on chromosome 9 (approximately 44 Mb). The near-zero marker-based additive variance indicates that, at this sample size, the trait carries no detectable common-variant signal, an outcome also observed for other quantitative disease traits in soybean association panels (Chang et al., 2016).

**Figure 6.**
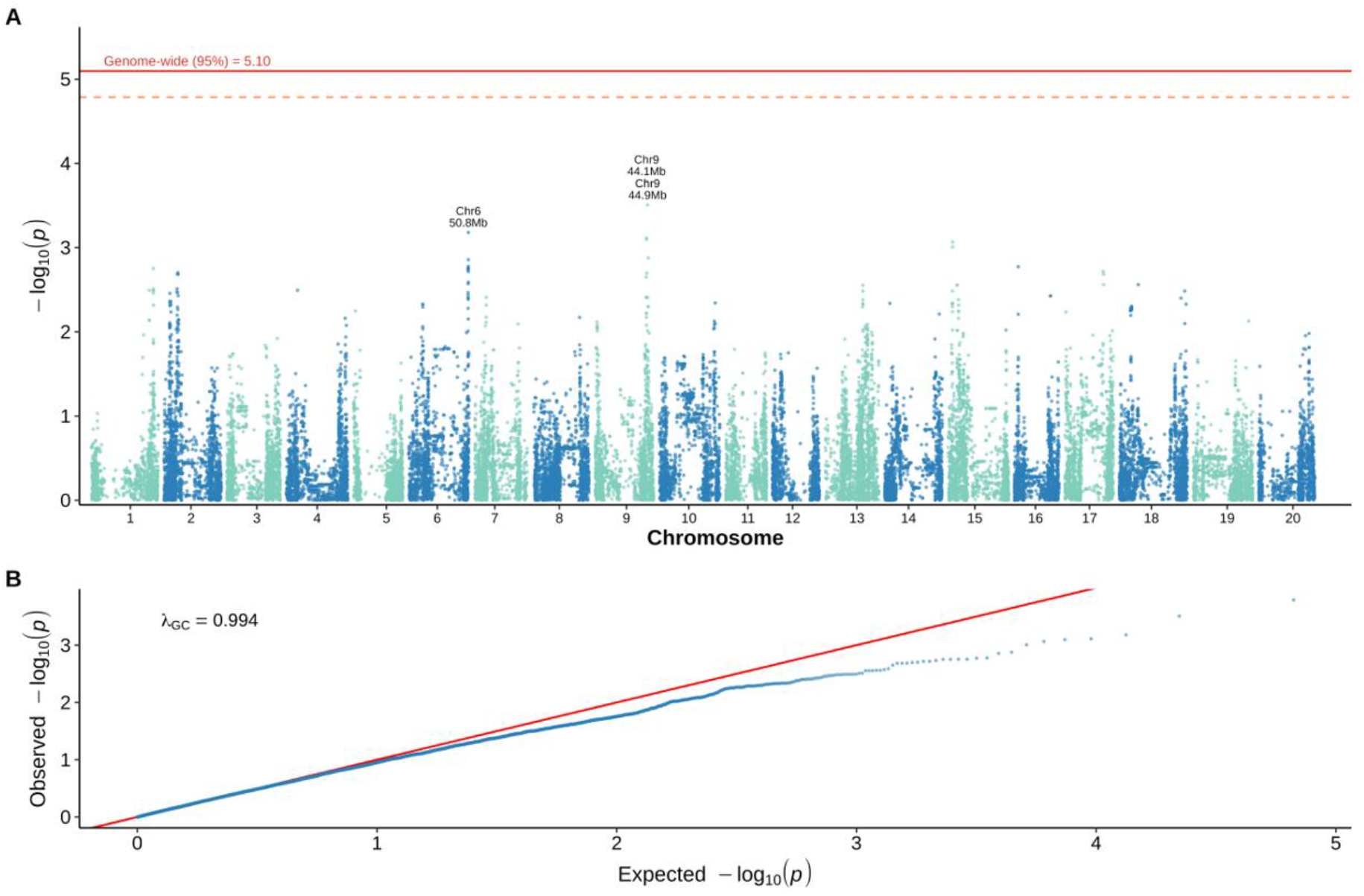
Genome-wide association analysis of the priming effect in the diversity panel (135 accessions, public SoySNP50K genotypes). (A) Manhattan plot with the permutation-derived significance (95th percentile) and suggestive (90th percentile) thresholds. (B) Quantile-quantile plot with a well-controlled genomic inflation factor (lambda_GC = 0.99). Fig. 6 alt text. Genome-wide association results for the diversity panel using public SoySNP50K markers. Panel A is a Manhattan plot with the permutation threshold marked and no marker exceeding it. Panel B is a quantile-quantile plot closely following the expected diagonal, indicating a well-calibrated analysis.

Cross-population meta-analysis. Combining the two GWAS at the level of shared 500-kb genomic windows (1,430 windows genotyped in both populations) revealed no region corroborated across populations: window-level signals were uncorrelated between the RIL and diversity panel (Pearson r = 0.02, P = 0.40), no window reached significance after false-discovery-rate correction, and no window showed elevated signal (-log10 P > 2) in both populations simultaneously (Supplementary Fig. S11) (Benjamini and Hochberg, 1995). The single RIL candidate region that could be directly tested on shared markers (chromosome 16, near GmNPR1-2 and GmWRKY72) showed no support in the panel. The convergence of three independent, well-powered analyses on the same negative result constitutes strong evidence that AHL-priming responsiveness lacks a common-variant, large-effect basis detectable at these sample sizes.

### 3.5 Exploratory RIL loci coincide with isoflavonoid and Defense-signaling candidate genes

Although no marker reached genome-wide significance, the strongest exploratory RIL signals recurred on chromosomes 5 and 16 and coincided with biologically relevant candidates (Supplementary Fig. S8). The chromosome-5 peak lay near a basic-leucine-zipper transcription factor gene of the isoflavonoid-regulatory class, the pathway that produces glyceollin (Akashi et al., 2009; Lin et al., 2024). The chromosome-16 peak coincided with GmNPR1-2, an ortholog of the systemic acquired resistance master regulator NPR1 (Withers and Dong, 2016; Backer et al., 2019), and with GmWRKY72, recently characterized as a direct transcriptional repressor of glyceollin biosynthesis (Lin et al., 2024). Grouping RILs by parental allele indicated allelic effects consistent with these candidates: at chromosome 5, Primus-type lines showed higher fold induction than Sigalia-type lines (P = 0.043), and at chromosome 16, the parental alleles differed in priming effect (P = 0.011) (Figure 7, Table 1). Given the null genome-wide result and the absence of cross-population support, these candidates are presented as hypotheses for future validation rather than as confirmed causal genes.

**Figure 7.**
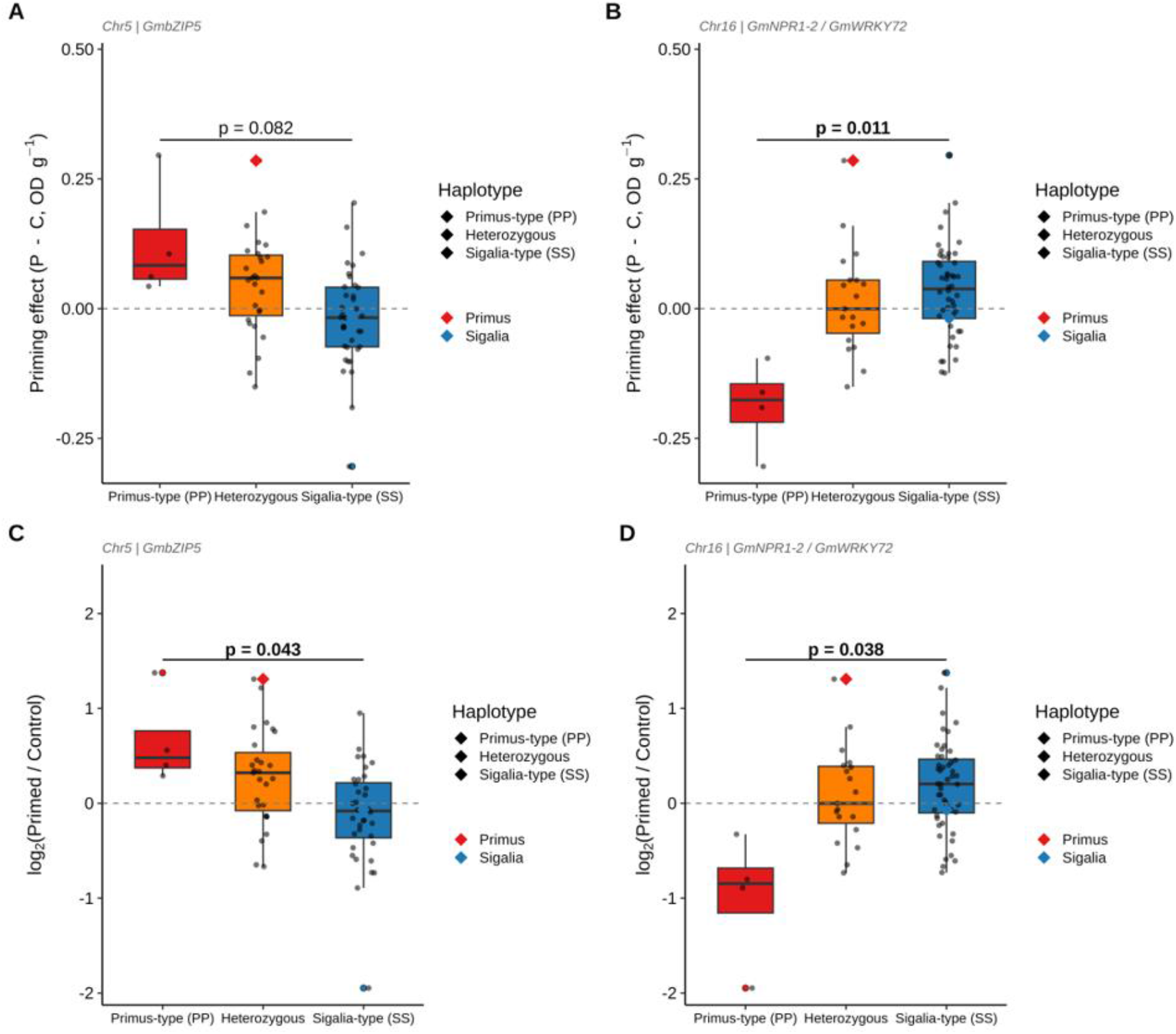
Allelic effects at the two leading exploratory loci in the RIL population. Priming effect (P minus C; A, B) and log2 fold induction (C, D) for RILs grouped by allele at the chromosome-5 (A, C) and chromosome-16 (B, D) peak SNPs. Groups are Primus-type homozygous, heterozygous, and Sigalia-type homozygous; parents are indicated. P-values are from t-tests comparing the two homozygous groups. Fig. 7 alt text. Box plots of the priming effect and of log-transformed fold induction for recombinant inbred lines grouped by parental allele at the two leading exploratory loci on chromosomes 5 and 16. Differences between allele groups are small and overlapping.

**Table 1.**
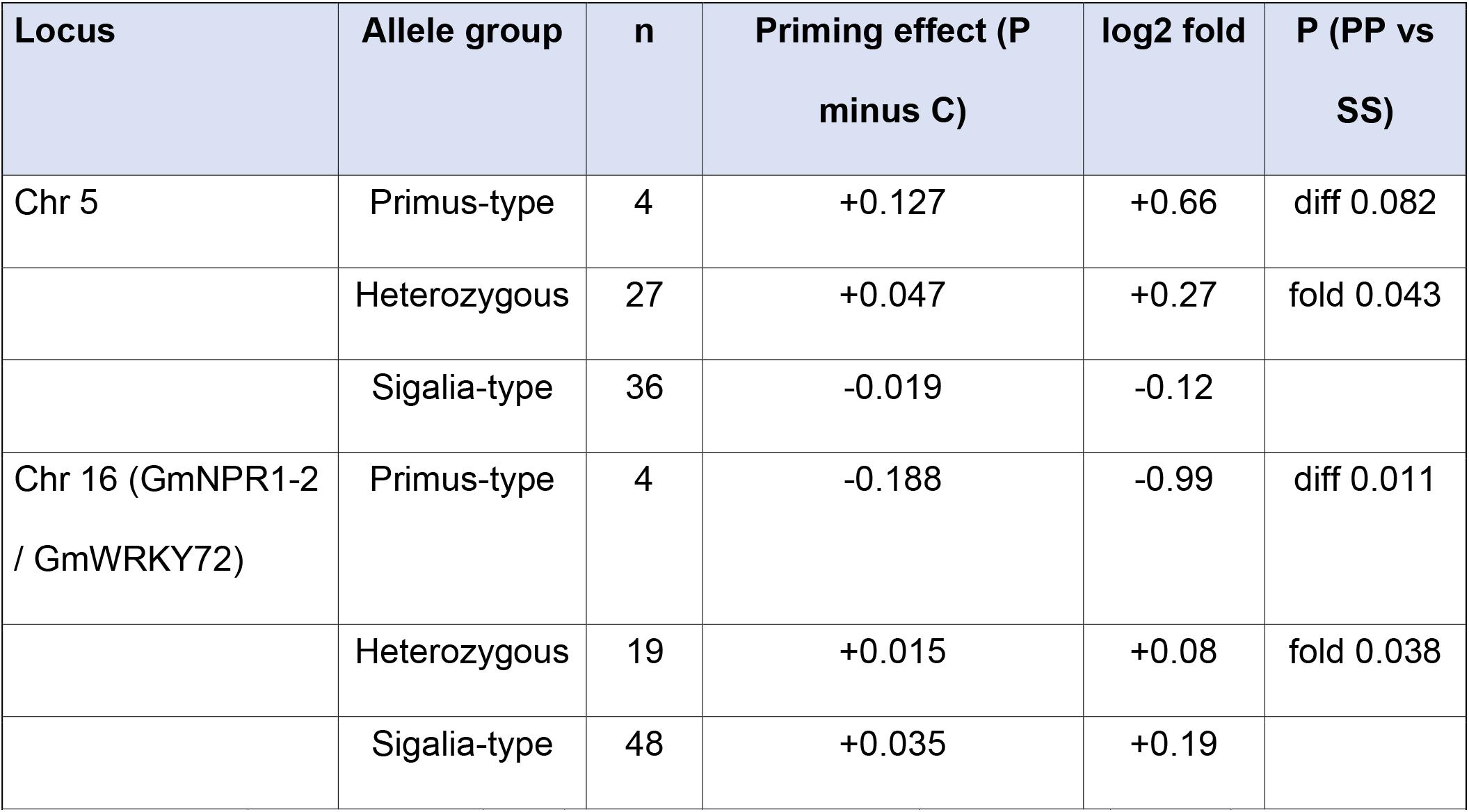
Allelic effects at the two leading exploratory loci on priming responsiveness in the Sigalia × Primus RIL population. Values are group means for the priming effect (P minus C, OD g⁻¹) and log2 fold induction; P-values compare Primus-type versus Sigalia-type homozygous groups (t-test).

**Table 2.**
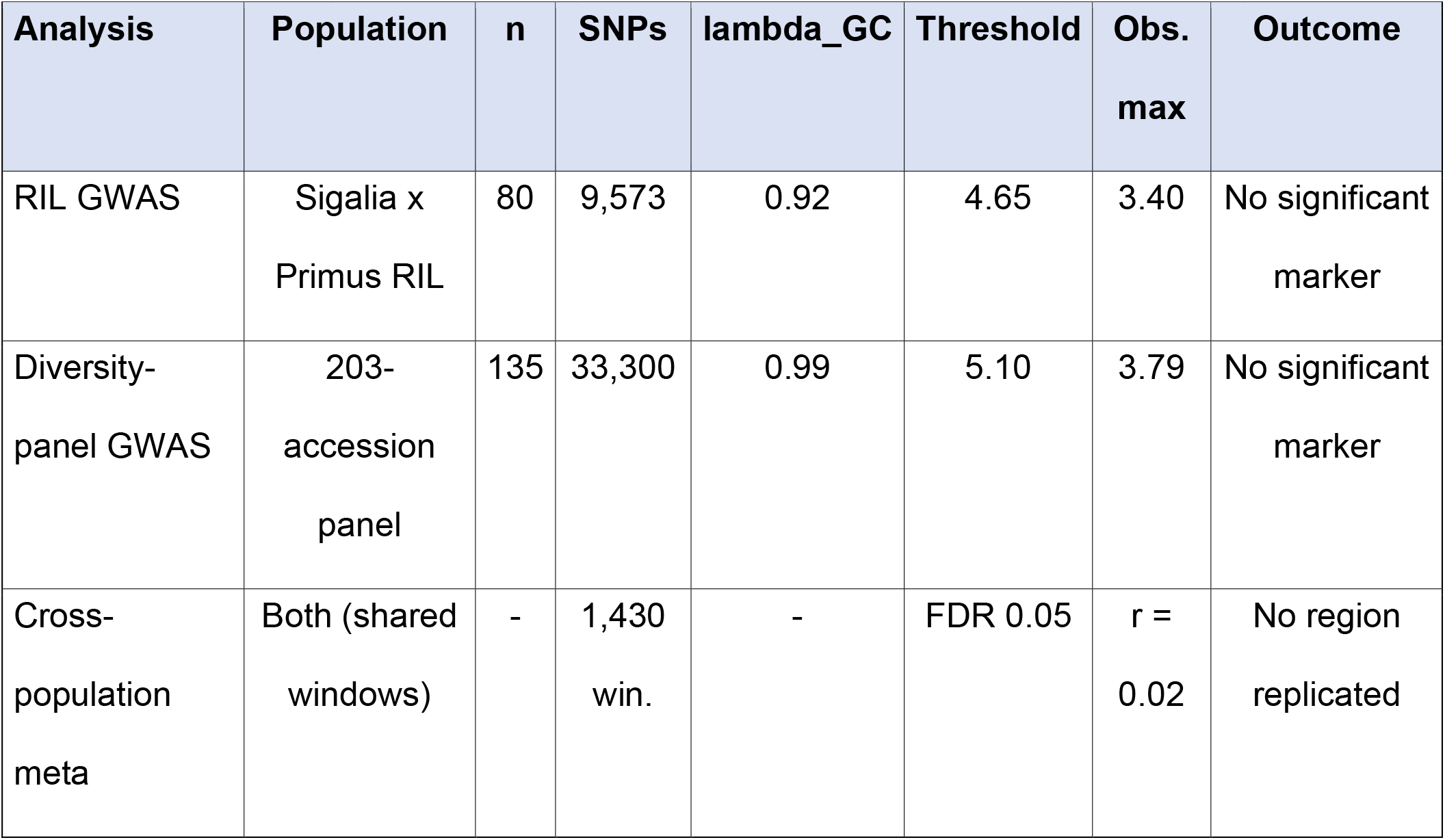
Summary of the three genetic analyses of the priming effect, illustrating convergent, well-powered null results across populations and marker platforms.

## 4. Discussion

This study establishes that the ability of soybean to translate the bacterial quorum-sensing signal oxo-C14-HSL into primed Defense is a genotype-dependent property of the plant. Both the oxo-C14-HSL-producing bacterium E. meliloti ExpR+ and the purified oxo-C14-HSL molecule suppressed nematode invasion in the responsive cultivar Primus but not in the weakly responsive cultivar Sigalia. Because the pure signal reproduced the contrast, the difference lies in the plant response to the AHL signal rather than in the bacterial colonization, consistent with reports that plant genotypes and species differ in their perception of and response to specific AHL molecules (Ortíz-Castro et al., 2008; Schenk and Schikora, 2014; Hernández-Reyes et al., 2014). Glyceollin accumulated in a priming- and challenge-dependent manner specifically in Primus, providing a mechanistic link between AHL perception and effective Defense (Adss et al., 2021). To our knowledge, this is the first demonstration in soybean that the capacity to convert a perceived AHL signal into primed phytoalexin Defense is under host genetic control, extending to a legume a principle of genotype-dependent AHL responsiveness previously established in barley (Shrestha et al., 2019; Wehner et al., 2019).

The glyceollin data exemplify priming as sensitization rather than constitutive activation: Primus did not over-accumulate glyceollin upon priming alone but responded strongly when priming was followed by challenge, the hallmark of a primed state in which Defenses are potentiated for rapid deployment at minimal standing cost (Conrath et al., 2015; Martínez-Medina et al., 2016; Mauch-Mani et al., 2017). The comparatively flat response of Sigalia identifies a genotype that perceives the priming stimulus, as shown by its response to the purified molecule, yet fails to transduce it effectively into a potentiated, phytoalexin-based Defense output. The genotype difference therefore resides in signal transduction and inducible-Defense capacity rather than in signal perception itself.

### 4.1 Candidate mechanisms linking AHL priming to glyceollin-based Defense

Several mechanistically coherent routes could explain the genotype-dependent coupling of AHL perception to glyceollin output. First, canonical AHL-priming in Arabidopsis and barley proceeds through augmented and prolonged activation of MPK3 and MPK6 and potentiated transcription of WRKY-type Defense regulators, downstream of the salicylic acid and oxylipin pathways (Schikora et al., 2011; Schenk et al., 2014; Schenk and Schikora, 2014). Because these kinases and transcription factors gate the amplitude of secondary-metabolite responses, genotypic variation in their activation could directly translate into the differential glyceollin induction we observed. Second, glyceollin biosynthesis itself is inducible and tightly controlled: it requires multiple enzymatic steps from daidzein and is governed by a balance of transcriptional activators, including GmNAC42-1 and GmMYB29A2, and repressors, notably GmWRKY72 (Graham et al., 2007; Jahan et al., 2020; Lin et al., 2024). GmWRKY72 directly binds glyceollin biosynthetic gene promoters and interacts with the activators GmNAC42-1 and GmMYB29A2 to restrain phytoalexin output (Lin et al., 2024); allelic or regulatory variation that relaxes this repression in Primus, or sustains it in Sigalia, offers a parsimonious explanation for the contrasting responsiveness. Third, effective priming depends on rapid flux through the isoflavonoid branch of the phenylpropanoid pathway, and the committed prenylation and cyclization steps catalyzed by GmG4DT and the recently identified P450 cyclases determine the rate of glyceollin isomer formation (Akashi et al., 2009; Xie et al., 2025). Genotypic differences in the inducibility of these committed enzymes could set the ceiling on how quickly a primed plant converts precursors into active phytoalexins in response to nematode challenge. Finally, the recently characterized AHL-priming protein ALI1, indispensable for oxo-C14-HSL-dependent priming in Arabidopsis, raises the possibility that variation in an upstream perception or relay component underlies the whole-plant contrast (Shrestha et al., 2022). These mechanisms are not mutually exclusive and disentangling them will require targeted expression and functional analyses in Primus and Sigalia.

Phenotyping across two independent populations, a biparental RIL population and an unrelated 203-accession diversity panel, showed that priming responsiveness is not a binary property of two cultivars but a continuously distributed, quantitative trait. Transgressive segregation in the RIL population and the broad distribution in the diversity panel, where numerous accessions exceeded Primus, indicate that favorable alleles are dispersed across germplasm and can, in principle, be combined beyond the level of either reference parent, an encouraging prospect for breeding (Hahn and Würschum, 2014; Wilkinson et al., 2019).

We deliberately dissected the genetic architecture within a power-aware, triangulated framework, and we regard the resulting convergence as a principal strength of the study rather than a limitation. Three independent analyses, a RIL GWAS, an orthogonal diversity-panel GWAS built on denser public markers and an entirely different set of accessions, and a cross-population meta-analysis, each returned a well-controlled null (lambda_GC = 0.92 and 0.99; no marker surpassing permutation-based significance; no region corroborated across populations). When three analyses that differ in population, marker platform, and statistical model converge on the same negative result, the most parsimonious interpretation is not that all three were coincidentally underpowered, but that the trait genuinely lacks a common-variant, large-effect basis (Churchill and Doerge, 1994; Korte and Farlow, 2013). This is reinforced by the low cross-experiment repeatability of the priming phenotype (r approximately 0.17): an inducible, environment-sensitive Defense trait of this kind offers a limited heritable signal to association mapping, and the honest, defensible conclusion is a power-informed null rather than an emphasis on sub-threshold peaks.

We nonetheless report exploratory RIL associations on chromosomes 5 and 16 because they recur across models and traits and coincide with biologically compelling candidates in the isoflavonoid-regulatory and systemic-resistance-signaling networks, including a locus near GmNPR1-2 and the glyceollin repressor GmWRKY72 (Withers and Dong, 2016; Lin et al., 2024). The direction of allelic effects at these loci is consistent with the biology of priming, since relaxation of WRKY72-mediated repression would be expected to raise the amplitude of the primed glyceollin response (Lin et al., 2024). However, these loci did not reach significance, and the one region testable across platforms did not replicate in the diversity panel; the candidates are therefore hypotheses for functional validation rather than established causal genes. Reporting candidates transparently while declining to over-interpret them is central to the defensibility of our conclusions.

Several boundaries define the scope of our inferences. The low phenotype repeatability argues that improved phenotyping precision, through greater replication, tighter environmental control, and complementary molecular read-outs of the primed state, such as Defense-gene expression, would yield more than added markers or samples alone. The RIL population is biparental and modest in size; the diversity-panel GWAS, though better powered and better calibrated, was necessarily restricted to the 135 accessions with public SoySNP50K genotypes, so the parental cultivars and several European and Asian accessions could not be included on the shared marker set. Genotyping these remaining accessions on a common platform would enable the parents to be anchored within the panel and would allow every candidate region to be tested across populations on identical markers, the definitive form of the cross-population analysis. Finally, a common-variant null does not exclude a polygenic architecture of many small effects, rare variants of larger effect, or a predominant role for the environment; distinguishing these will require larger, repeatedly phenotyped panels and complementary approaches such as expression profiling and allele-specific comparisons in Primus and Sigalia (Akashi et al., 2009; Lin et al., 2024).

In summary, the capacity to translate a bacterial quorum-sensing signal into primed phytoalexin Defense is a heritable, quantitatively varying property of soybean, mechanistically linked to inducible glyceollin accumulation and broadly distributed across germplasm. By establishing this phenotype, demonstrating its quantitative inheritance in two independent populations, and characterizing its genetic architecture through a rigorous, triangulated analysis, our study provides both a defined quantitative phenotype and a defensible genetic framework for dissecting natural variation in inducible Defense, and ultimately for improving microbiome-primed resistance in soybean.

## Supplementary data

The following supplementary data are available at JXB online.

*Fig. S1. Nematode density in Sigalia primed with E. meliloti attM, E. meliloti ExpR+, or control (no significant difference)*.

*Fig. S2. Glyceollin accumulation in control versus primed cotyledons across the RIL population*.

*Fig. S3. RIL priming responsiveness expressed as fold induction (P/C)*.

*Fig. S4. RIL GWAS phenotype distribution and marker properties*.

*Fig. S5. RIL population structure: PCA, kinship, and LD decay*.

*Fig. S6. Null distribution of maximum -log10 P from 1,000 permutations (RIL)*.

*Fig. S7. RIL general-linear-model Manhattan and QQ plots*.

*Fig. S8. Locus-zoom plots of the RIL chromosome-5 and chromosome-16 candidate regions*.

*Fig. S9. RIL fold-induction GWAS Manhattan and QQ plots*.

*Fig. S10. Diversity-panel population structure (PCA)*.

*Fig. S11. Cross-population comparison of window-level signals and region-level meta-analysis*.

*Table S1. RIL marker quality-control summary*.

***Table S2. Complete list of the 207 diversity-panel accessions with registry codes, accession names, USDA Plant Introduction (PI) numbers, and public SoySNP50K genotype availability.***

*Table S3. Permutation-derived significance thresholds for both GWAS*.

***Table S4. Diversity-panel marker availability.***

## Acknowledgements

The authors thank Prof. Dr. Adam Schikora for providing the bacterial strains and sharing protocols. We gratefully acknowledge the USDA-ARS Soybean Genomics and Improvement Laboratory and SoyBase for public access to the SoySNP50K genotype data.

## Author contributions

H.H. and A.E. conceptualized and designed the study. S.A. and A.E. performed the experiments. S.A. and A.E. analyzed the genomic and phenotyping data. S.A. wrote the first draft of the manuscript and designed the figures. V.H. provided the seed material, genetic features, and passport information for the accessions. S.A., A.E., L.B., V.H., and H.H. reviewed and edited the manuscript and approved the final version for submission. All authors have read and agreed to the published version of the manuscript.

## Data availability

The USDA Plant Introduction (PI) accession numbers for all diversity-panel accessions are listed in Supplementary Table S2, and the genotypes of the PI accessions are publicly available from the SoyBase SoySNP50K resource (https://soybase.org). The raw genotyping-by-sequencing reads for the RIL population have been deposited in the European Nucleotide Archive (ENA), and the accession number will be provided prior to publication. Phenotype data and processed marker matrices are available from the corresponding author upon reasonable request.

